# Elevated *MCOLN1* Expression in p53-Deficient Bladder Cancer is Necessary for Oncogene-Induced Cell proliferation, Inflammation, and Invasion

**DOI:** 10.1101/2020.07.08.193862

**Authors:** Jewon Jung, Han Liao, Hong Liang, John F. Hancock, Catherine Denicourt, Kartik Venkatachalam

## Abstract

Inhibition of the endolysosomal cation channel, TRPML1, which is encoded by *MCOLN1*, deters the proliferation of cancer cells with augmented TFEB activity. Here, we report that the tumor suppressor, p53, antagonizes TFEB-driven *MCOLN1* expression in bladder cancer. Not only was the constitutive loss of p53 in bladder cancer cells associated with higher *MCOLN1* mRNA, knockdown of *TP53* in lines with wild type alleles of the tumor suppressor increased *MCOLN1* expression. Elevated TRPML1 abundance in p53-deficient cancer cells, although not sufficient for bolstering proliferation, was necessary for the effects of oncogenic HRAS on cell division, cytokine production, and invasion. These data demonstrate that hyperactivation of the TFEB– *MCOLN1* transcriptional axis in urothelial cells lacking p53 permits tumorigenesis stemming from *HRAS* mutations. Furthermore, the insight that loss of p53 predicts addiction to TRPML1 informs an actionable therapeutic strategy for bladder cancer.

## Introduction

By coordinating the synthesis and breakdown of macromolecules with signaling events that regulate metabolism, endolysosomes play vital roles in cellular homeostasis (Xu and Ren, 2015). While insufficient endolysosomal function is associated with inborn errors of metabolism, lysosomal storage, and degenerative diseases, increased biogenesis of these organelles is a recurring theme in cancers (Davidson and Vander Heiden, 2017). Several studies have described roles for the endolysosomal cation channel, TRPML1, in different malignancies (Hu et al., 2019; Jung et al., 2019; Kasitinon et al., 2019; Xu et al., 2019; Yin et al., 2019). A common theme in these studies is that TRPML1 is necessary for malignancy promoting signaling cascades involving RAS–MAPK and mTORC1. For instance, *HRAS*-driven cancers exhibit increased TRPML1 abundance and activity, which underlies endolysosomal exocytosis necessary for recycling cholesterol-rich membranes and oncogenic HRAS to the plasma membrane (Jung and Venkatachalam, 2019; Jung et al., 2019). As a result, TRPML1 inhibition or *MCOLN1* knockdown disrupts ERK activation and cell proliferation in HRAS-driven cancer cells. TRPML1-dependent vesicle exocytosis is also necessary for the release of lysosomal ATP, which in the case of triple negative breast cancers, stimulates cell migration and metastases (Xu et al., 2019). In other cancers, TRPML1 promotes untrammeled proliferation and tumor growth via mTORC1 (Hu et al., 2019; Xu et al., 2019).

These studies make a compelling argument for understanding the pathways that govern *MCOLN1* expression in cancers. Of major significance here are TFEB, TFE3, and MITF — transcription factors that induce the expression of *MCOLN1* and other endolysosomal genes harboring ‘coordinated lysosomal expression and regulation’ (CLEAR) motifs (Martina et al., 2014; Palmieri et al., 2011; Ploper et al., 2015; Sardiello et al., 2009; Settembre et al., 2011, 2012). Signaling events that influence the nucleocytoplasmic distribution of these transcription factors function as *de facto* regulators of endolysosomal biogenesis (Martina et al., 2012; Medina et al., 2015; Palmieri et al., 2017; Roczniak-Ferguson et al., 2012; Settembre et al., 2011). Several cancers are characterized by either the constitutive localization of TFEB/TFE3/MITF in cell nuclei, or by amplifications that upregulate their expression (Argani, 2015; Blessing et al., 2017; Calcagnì et al., 2016; Kauffman et al., 2014; Kundu et al., 2018; Li et al., 2019; Perera et al., 2015, 2019; Ploper et al., 2015). The attendant potentiation of endolysosomal biogenesis supports cancer cell survival, proliferation, and metastases. Since inhibition of TRPML1 arrests proliferation or induces cell death in cancers with elevated expression of the CLEAR network genes (Hu et al., 2019; Jung and Venkatachalam, 2019; Jung et al., 2019), tumors with hyperactivated TFEB/TFE3/MITF signaling are addicted to TRPML1.

Despite all this progress, many questions persist. For instance, we still do not fully understand how the CLEAR network and *MCOLN1* expression are pathologically augmented in cancer. In healthy cells, feedback mechanisms keep endolysosomal biogenesis in check. For instance, mTORC1 suppresses endolysosomal biogenesis by driving phosphorylation and cytosolic retention of TFEB and TFE3 (Martina et al., 2012, 2014; Roczniak-Ferguson et al., 2012; Wang et al., 2015), which is why high mTORC1 activity is correlated with repression of the CLEAR network. In cancers, however, these axes are often dysregulated — a notion exemplified by KRAS-driven pancreatic cancer cells, which exhibit hyperactivation of *both* mTORC1 and TFEB-driven gene expression due to failure of the mutually inhibitory mechanisms linking these entities (Perera et al., 2015). In addition, genomic landscapes of cancer cells encourage qualitative shifts in the regulatory pathways that control endolysosomal biogenesis. Signaling that normally curtails endolysosomal gene expression morphs into drivers of vesicular biogenesis in cancer. For example, while the tumor suppressor, p53, activates TFEB and TFE3 in normal fibroblasts exposed to DNA damage, loss of p53 in cancers is also associated with the paradoxical activation of the TFEB/TFE3–endolysosome axis (Brady et al., 2018; Tasdemir et al., 2008a, 2008b; Zhang et al., 2017). The lack of clear insights into the regulation of TFEB/TFE3-driven endolysosomal biogenesis hinders our ability to exploit TRPML1 addiction as a therapeutic strategy. To fill the gaps in knowledge, we surveyed *MCOLN1* expression in different cancers using the cancer genome atlas (TCGA) (Cancer Genome Atlas Research Network, 2014). This analysis prompted a focus on bladder carcinoma (BLCA) (Robertson et al., 2017), in which primary tumors exhibited significant elevations in *MCOLN1* expression. Further investigation revealed a role for p53 in repressing TFEB-driven *MCOLN1* expression. Therefore, loss of p53 augmented TRPML1 abundance, which in turn fostered cell proliferation, inflammation, and invasion stemming from oncogenic HRAS. Our study uncovers a novel axis by which *MCOLN1* expression is regulated, and suggests that TRPML1 inhibitors could mitigate tumorigenesis in p53-deficient bladder cancers.

## Results

### *MCOLN1* expression was inversely correlated with p53 targets in bladder cancer

By comparing gene expression in tumors relative to matched normal tissues, we found that *MCOLN1* expression was elevated in BLCA (Figure 1A and Supplementary Table1), which prompted us to select this disease as a suitable model for the identification of cancer-related pathways that depend on increased *MCOLN1* expression. We also reasoned that ontologies of genes whose transcription correlates with *MCOLN1* would reveal the pathways that depend on precise calibration of *MCOLN1* expression. In BLCA, *MCOLN1* exhibited significant positive and negative correlation with 4737 and 3611 genes, respectively (Figure 1B and Supplementary Table 2). Targeted gene set enrichment analysis (GSEA) (Subramanian et al., 2005) using the correlation coefficients revealed the expected enrichment of CLEAR targets in genes that are positively correlated with *MCOLN1* (Figure 1C) (Palmieri et al., 2011). Likewise, upon probing the correlation coefficients against MSigDB, we found that genes positively correlated with *MCOLN1* exhibited enrichment of modules related to lysosomes and lytic vacuoles, endocytosis and phagocytosis, and vesicular exocytosis and secretion (Figures 1D and S1). In addition, *MCOLN1* expression was positively correlated with genes that are upregulated during UV-induced DNA repair, and negatively correlated with p53 target genes and genes that are repressed during UV-induced DNA repair (Figures 1D and S1).

**FIGURE 1.**
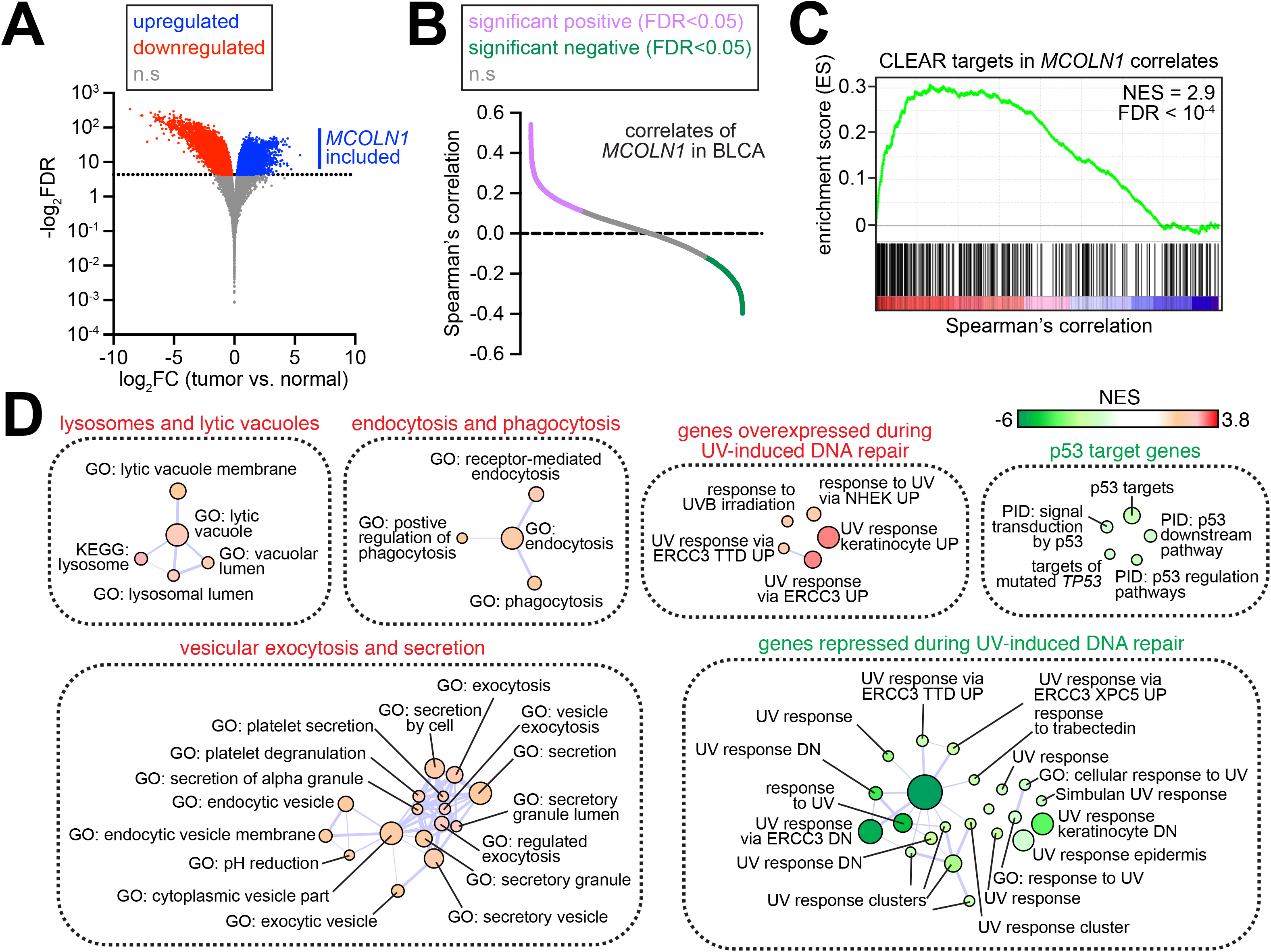
Pathways correlated with *MCOLN1* expression in BLCA. **(A)** Volcano plot showing DEGs in BLCA primary tumors relative to matched normal tissues. Genes that were significantly (FDR < 0.05) upregulated and downregulated in tumors are depicted by blue and red dots, respectively. *MCOLN1* belonged to the set of significantly upregulated genes. Gray dots represent genes whose expression was not significantly altered in tumors. **(B)** Plot showing correlation of 20,177 genes with *MCOLN1*. Values on the Y-axis represent Spearman’s correlation. Genes that showed significant (FDR < 0.05) positive and negative correlation with *MCOLN1* are represented by magenta and green dots, respectively. Gray dots represent genes that were not significantly correlated with *MCOLN1*. **(C)** GSEA shows enrichment for CLEAR targets in genes that showed significant positive correlation with *MCOLN1*. **(D)** Functional annotation of genes that were significantly correlated with *MCOLN1* in BLCA. All the genes that were significantly correlated with *MCOLN1* were subjected to GSEA. Sets with GSEA FDR < 0.05 are shown as nodes. Node color represents NES from GSEA. Node size represents number of genes that make up that set. Thickness of the lines connecting the nodes indicates extent of overlap. Unconnected nodes have no overlap. Abbreviations: FC, fold change; NES, normalized enrichment score.

### *MCOLN1* expression was elevated in primary BLCA tumors with *TP53* mutations

Next, we applied the principles of information theory (Shannon, 1948) to determine whether mutations in any of the 722 genes belonging to the Cancer Gene Census (CGC; https://cancer.sanger.ac.uk/census; Supplementary Table 3) correlated with *MCOLN1* expression in BLCA. We sought to identify those genes that were mutated in tumors with either high or low *MCOLN1* expression, such that partitioning the tumors on the basis of *MCOLN1* expression resulted in a decrease in the apparent stochasticity of the mutations (i.e. an increase in ‘Shannon information’, see *Materials and Methods*). We found that 145 genes exhibited significant information gain upon partitioning the tumors on the basis of *MCOLN1* expression (Figure 2A and Supplementary Table 4). Of these genes, only four (*TP53, KDM6A, ARID1A, RB1*) were mutated in ≥15 tumors (Figure 2A). To validate the relationship between *MCOLN1* expression and mutations in these four genes, we performed targeted GSEA by ranking tumors on the basis of *MCOLN1* expression. We found that mutations in *TP53* (*TP53^mut^*) and *RB1* (*RB1^mut^*) were significantly enriched in tumors with highest *MCOLN1* expression, whereas wild type alleles of both genes (*TP53^wt^* and *RB1^wt^*) were enriched in tumors with lowest *MCOLN1* expression (Figure 2B). In agreement, pairwise comparison of normalized *MCOLN1* FPKM revealed significantly higher values in *TP53^mut^* and *RB1^mut^* in comparison to *TP53^wt^* and *RB1^wt^* tumors, respectively (Figure 2C). Mutations in *ARID1A* (*ARID1A^mut^*) or *KDM6A* (*KDM6A^mut^*), however, were not enriched in tumors with either the highest or lowest *MCOLN1* expression (Figure S2A). Neither was *MCOLN1* expression significantly altered in *ARID1A^mut^* or *KDM6A^mut^* tumors relative to tumors with wild type alleles of these two genes (*ARID1A^WT^* or *KDM6A^WT^*, respectively) (Figure 2C). Together, these data point to elevated *MCOLN1* expression in tumors harboring *TP53* and *RB1* mutations. Since tumors with *RB1* mutations belonged to the larger set with *TP53* mutations (Figure S3), we conclude that tumors harboring *TP53* mutations constitute the broadest set of samples with elevated *MCOLN1* expression.

**FIGURE 2.**
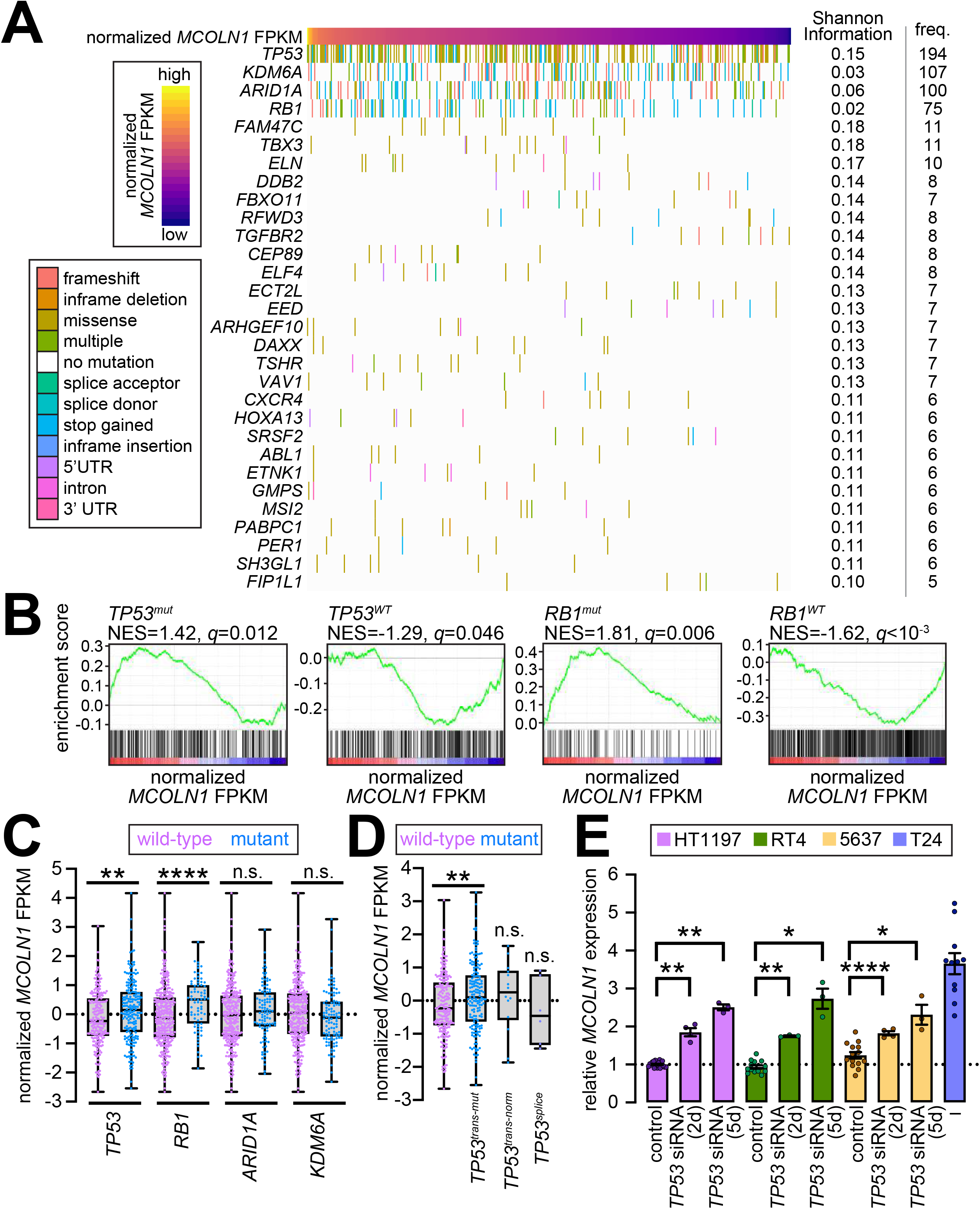
*MCOLN1* expression is elevated in *TP53^mut^* BLCA tumors. **(A)** Waterfall plot showing mutations in 30 genes with significant information gain. *MCOLN1* expression is depicted as median normalized FPKM values at the top. Bars mark tumors harboring mutations in respective genes, and bar colors represent the type of mutation. On the right, Shannon information associated with partitioning tumors into three groups based on *MCOLN1* expression are show; and ‘freq.’ depicts number of tumors in the BLCA dataset that exhibit mutations in the respective genes. For all the genes shown, *Q* < 0.0001. **(B)** GSEA traces show enrichment for indicated mutations in tumors ranked on the basis of *MCOLN1* expression (represented as median normalized FPKM values). **(C-D)** Box plots showing *MCOLN1* expression (represented as median normalized FPKM values) in BLCA sorted on the basis of the presence or absence of mutations in the indicated genes. Each dot represents the values from a different tumor and lines in the box plots represent median values. In **(C)**, **, *P* < 0.01, t-test; ****, *P* < 0.0001, Mann-Whitney tests. In **(D)**, **, *P* < 0.01, t-test. **(E)** Bar graph showing relative *MCOLN1* expression in the indicated bladder cancer cell lines treated with control siRNA or siRNA against *TP53* (75 nM) for either 2-days or 5-days (2d and 5d, respectively). For each cell type values were normalized to the control mean. Values for T24 cells were normalized to HT1197 mean. Dashed line indicated control mean. Circles represent independent biological repeats and the values shown represent mean ± SEM; *, *P* < 0.05; **, *P* < 0.01; ****, *P* < 0.0001, t-tests. Abbreviation: NES, normalized enrichment score; *q*, FDR value; n.s., not significant.

### *MCOLN1* expression was higher in BLCA tumors with *TP53* mutations that occur in the DNA binding domain

Approximately 47% of BLCA tumors in the TCGA dataset were *TP53^mut^*. With help of the IARC database (https://p53.iarc.fr/Default.aspx) (Bouaoun et al., 2016), we found that among the *TP53^mut^* BLCA tumors, ~86% harbored *TP53* mutations (*TP53^trans-mut^*) predicted to result in loss of transactivation function due to either nonsense mutations leading to premature termination or missense mutations in the DNA binding domain. Among the remainder, ~7% harbored *TP53* mutations (*TP53^trans-norm^*) that were either unlikely to affect transactivation or have remained unclassified. Another ~3% carried *TP53* mutations in splice acceptor or donor sites (*TP53^splice^*), and ~3% harbored multiple *TP53* mutations. Finally, one tumor harbored an inframe mutation. In sum, \majority of *TP53* mutations in BLCA were predicted to result in loss of p53 transactivation. Targeted GSEA revealed that *TP53^trans-mut^* mutations were significantly enriched in tumors with high *MCOLN1* expression (Figure S2B). In comparison to the values *inTP53^WT^* tumors, *MCOLN1* expression was significantly higher in *TP53^trans-mut^* tumors, but not in *TP53^trans-norm^* or *TP53^splice^* tumors (Figure 2D). Therefore, *MCOLN1* expression was elevated in tumors that harbored mutations in the DNA binding domain of p53.

### *MCOLN1* expression is elevated in p53-deficient bladder cancer cell lines

To determine whether the relationship between p53 and *MCOLN1* occurs in bladder cancer cells, we examined four different lines: HT1197, RT4, 5637, and T24 (Bubeník et al., 1973; Fogh, 1978; Rasheed et al., 1977; Rigby and Franks, 1970). As per the cancer cell line encyclopedia, HT1197 and RT4 are wild type for *TP53* (Supplementary Table 5). 5637 cells harbor a missense *TP53* mutation that is predicted to encode a transactivation-deficient (R280T) variant, and T24 cells are homozygous for a nonsense mutation in the *TP53* coding sequence that is predicted to yield a null allele (Supplementary Table 5). 5637 cells are also homozygous for a nonsense mutation in *RB1* (Supplementary Table 5). Using RT-PCR, we found that in comparison to the values in HT1197, RT4, or 5637 cells, *MCOLN1* expression was significantly higher in T24 cells (Figure 2E). These data indicate that loss of p53, but not Rb1, was associated with higher *MCOLN1* expression. In further agreement, *TP53* knockdown significantly elevated *MCOLN1* expression in HT1197, RT4, and 5637 cells (Figure 2E). Together, these data point to the sufficiency of p53 in repressing *MCOLN1* in bladder cancer cells.

Consistent with TFEB being a transcriptional activator of *MCOLN1* (Figure 3A) (Sardiello et al., 2009), knockdown of *TFEB* in T24 cells restored *MCOLN1* expression to control levels (Figure 3B). Furthermore, simultaneous knockdown of *TFEB* dampened the extent of *MCOLN1* induction stemming from *TP53* knockdown in RT4 cells (Figure 3C). Since TFEB reinforces its own expression (Figure 3A) (Jung et al., 2019; Sardiello et al., 2009), T24 cells also expressed higher levels of *TFEB* than did HT1197 and RT4 cells (Figure 3D). Likewise, knockdown of *TP53* was sufficient to significantly increase *TFEB* mRNA levels in both HT1197 and RT4 cells (Figure 3D). Together, these data are consistent with the notion that loss of p53 activates the TFEB–*MCOLN1* transcriptional axis.

**FIGURE 3.**
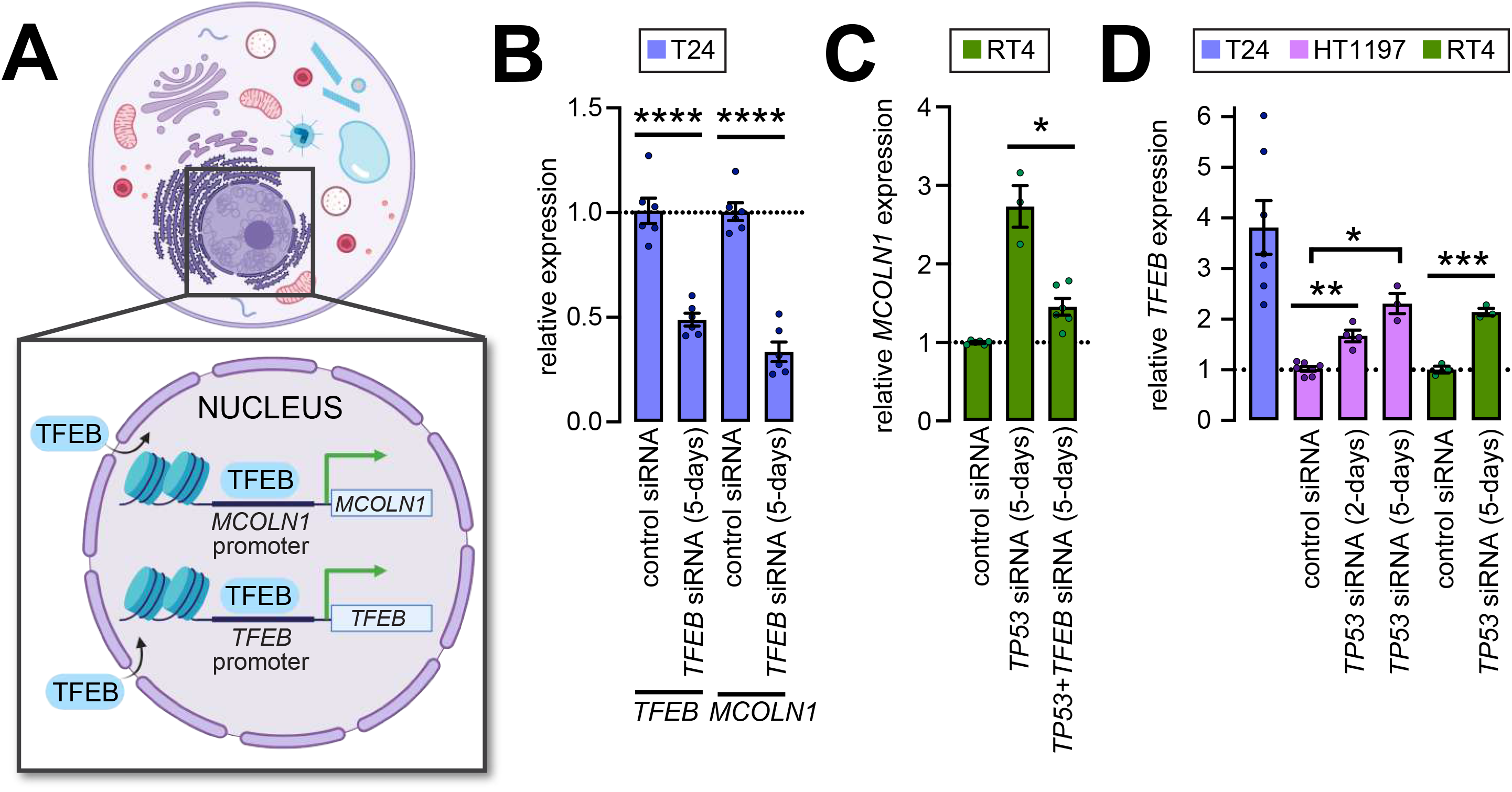
*MCOLN1* overexpression in the absence of p53 depends on TFEB. **(A)** Schematic showing that promoters of both *MCOLN1* and *TFEB* harbor CLEAR elements. Upon translocation into the nucleus, TFEB binds the promoters bearing CLEAR elements leading to the induction of the genes. **(B)** Relative *TFEB* and *MCOLN1* expression in T24 cells lines treated with control or 50 nM *TFEB* siRNA. All values were normalized to control mean. Circles represent independent biological repeats and the values shown represent mean ± SEM; ****, *P* < 0.0001, t-tests. **(C)** Relative *MCOLN1* expression in RT4 cells lines treated with control, 75 nM *TP53* siRNA, or 75 nM *TP53* siRNA and 50 nM *TFEB* siRNA. All values were normalized to control mean. Circles represent independent biological repeats and the values shown represent mean ± SEM; *, *P* < 0.05, t-test. **(D)** Relative *TFEB* expression in the indicated cells lines treated with nothing, control siRNA or 75 nM *TP53* siRNA. All values were normalized to mean in HT1197 cells treated with control siRNA. Circles represent independent biological repeats and the values shown represent mean ± SEM; *, *P* < 0.05, **, *P* < 0.01, ***, *P* < 0.001, t-tests with Bonferroni correction in case of samples used in multiple pairwise comparisons.

### Roles for TRPML1 in proliferation and prevention of cell cycle arrest in p53-deficient bladder cancer cells

Next, we identified the genes whose expression was significantly different in *TP53^mut^* tumors relative to those that were *TP53^wt^*. At FDR < 0.05, we identified 5659 differentially expressed genes (DEGs). *MCOLN1* belonged to the cohort of significantly upregulated genes in *TP53^mut^* tumors, whereas expressions of both *TP53* and *RB1* were significantly diminished (Figure 4A and Supplementary Table 6). GSEA identified several functional modules in the DEGs including those related to positive regulation of cell cycle progression and proliferation, and immunity and inflammation (Figure 4B and Supplementary Table 7). Additional upregulated modules were comprised of gene sets for cell adhesion, RNA processing, and response to zinc (Figure 4B and Supplementary Table 7). Downregulated modules involved those related to detoxification/drug metabolism via cytochrome p450, hormone signaling, lipase activity, ribosome and protein translation, and solute pumps (Figure 4B and Supplementary Table 7).

**FIGURE 4.**
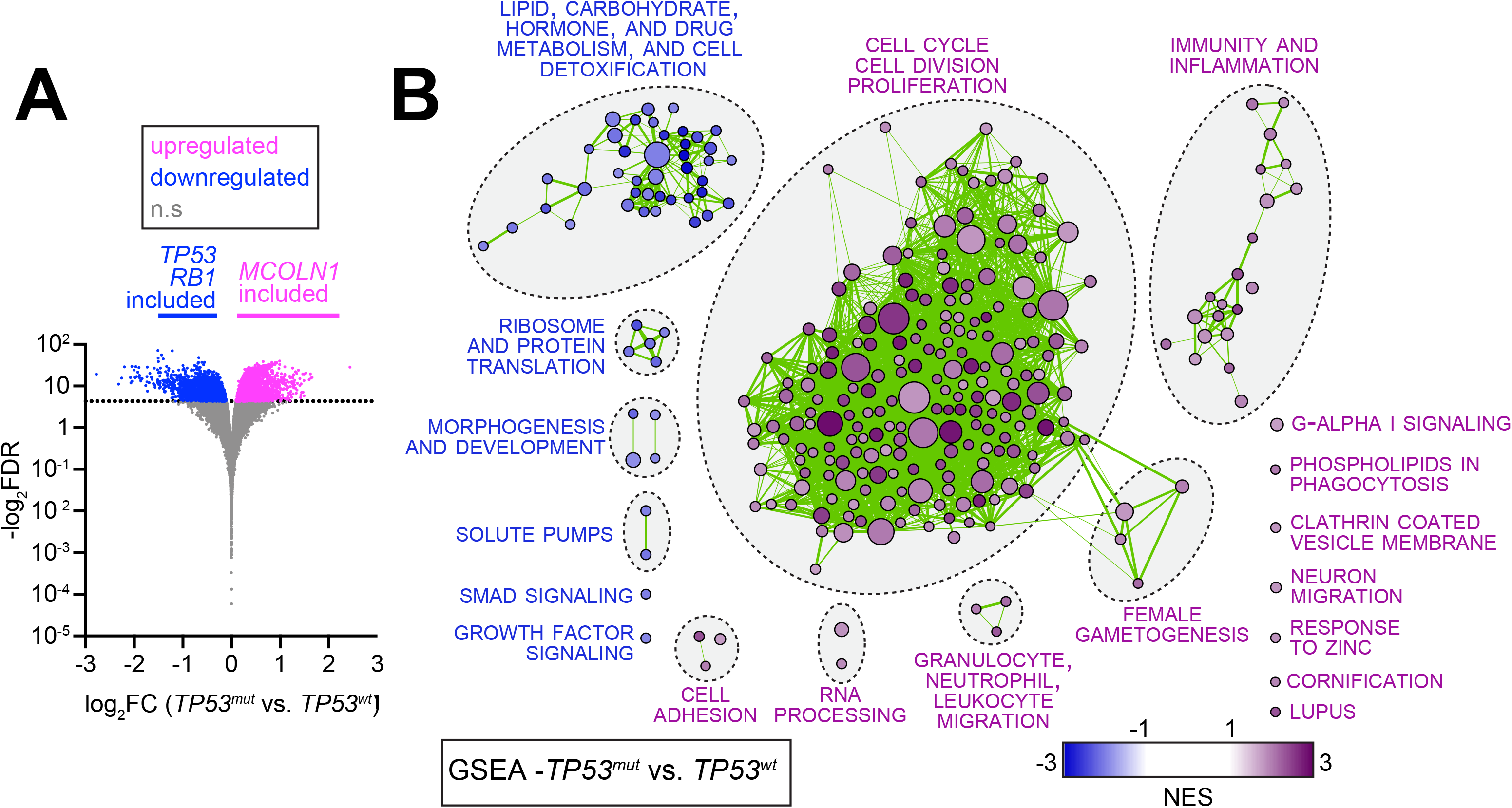
Transcriptional alterations in BLCA tumors harboring *TP53* mutations. **(A)** Volcano plot showing DEGs in *TP53^mut^* compared to *TP53^wt^* tumors. Genes that were significantly (FDR < 0.05) upregulated and downregulated in *TP53^mut^* tumors are depicted by magenta and blue dots, respectively. *MCOLN1* belonged to the set of genes upregulated genes, whereas *TP53* and *RB1* belonged to the set of downregulated genes. Gray dots represent genes whose expression was not significantly altered. **(B)** Functional annotation of DEGs in *TP53^mut^* relative to *TP53^wt^* BLCA tumors. All DEGs were subjected to GSEA. From the results of this analysis, only the gene sets with GSEA FDR < 0.05 are shown as nodes. Node color represents NES from GSEA. Node size represents number of genes that make up that set. Thickness of the lines connecting the nodes indicates extent of overlap. Unconnected nodes have no overlap. Abbreviations: FC, fold change.

Since modules related to cell cycle progression were upregulated in *TP53^mut^* tumors, we asked whether TRPML1 might be necessary for the proliferation of bladder cancer cells lacking p53. Using the MTT cell proliferation assay, we found that pharmacological inhibition of TRPML1 by application of ML-SI1 (Samie et al., 2013) for 2-days significantly attenuated T24 cell proliferation (Figure 5A). In contrast, HT1197 and RT4 cells were much less sensitive to TRPML1 inhibition (Figure 5A). Extending the duration of ML-SI1 treatment to 5-days led to even greater attenuation of T24 cell numbers, whereas the extended regimen did not further attenuate the proliferation of either HT1197 or RT4 cells (Figure 5A). To rule out the possibility that the anti-proliferative effects of ML-SI1 reflect a specific influence on the MTT assay, we assessed colony formation. ML-SI1 application led to significantly greater reduction in the number of colonies in T24 relative to HT1197 (Figure 5B). Furthermore, *MCOLN1* siRNA curtailed the proliferation to a significantly greater extent in T24 than in HT1197, RT4, or 5637 cells (Figure 4C). In agreement with our previous findings that this siRNA reduces *MCOLN1* expression by ~50% in T24 cells (Jung et al., 2019), *MCOLN1* knockdown in HT1197, RT4, and 5637 cells was also ~50% (Figure S1A). Therefore, the relative insensitivity of HT1197, RT4, and 5637 cells to *MCOLN1* siRNA was not due insufficient knockdown.

**FIGURE 5.**
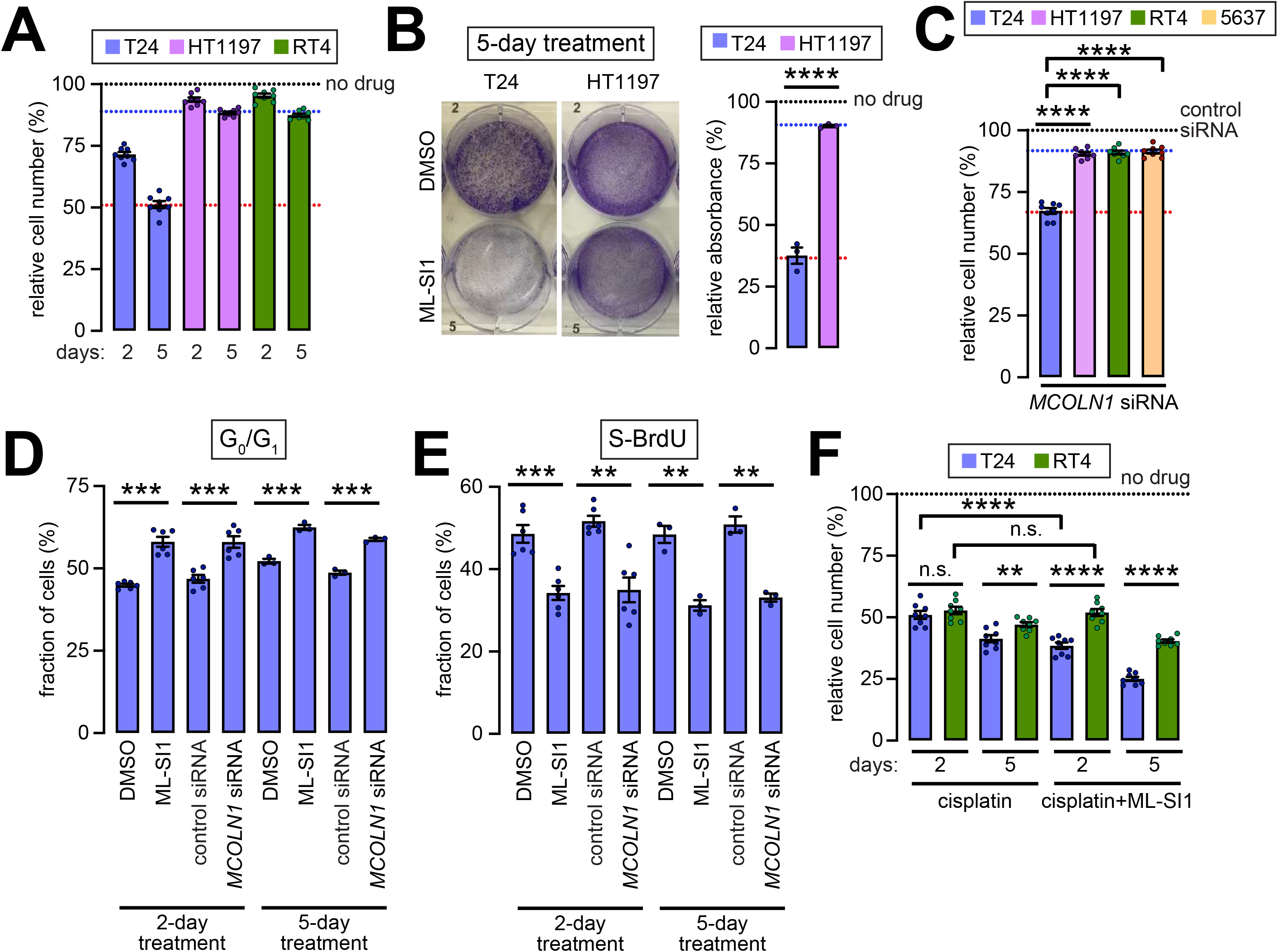
TRPML1 is required for the proliferation of T24, but not HT1197, RT4, or 5637 bladder cancer cells. **(A)** Bar graph showing relative number of the indicated cells assessed using the MTT cell proliferation assay. Numbers at the bottom represent duration of 10 μM ML-SI1 application in days. Black dashed line represents control values for each cell line in the absence of any drug treatment. Red dashed line represents values in T24 cell treated with ML-SI1 for 5-days. Blue dashed line represents values in HT1197 and RT4 cell treated with ML-SI1 for 5-days. Circles represent independent biological repeats and the values shown represent mean ± SEM. **(B)** *Left*, representative images of the indicated cell lines treated with DMSO or 10 μM ML-SI1 for 5-days. Cells grown on the dished were stained with crystal. *Right*, bar graph showing intensity of crystal violet staining in ML-SI1 treated cells normalized to the values in untreated cells. Black dashed line represents control values for each cell line in the absence of any drug treatment. Red dashed line represents values in T24 cell treated with ML-SI1 for 5-days. Blue dashed line represents values in HT1197 cells treated with ML-SI1 for 5-days. Circles represent independent biological repeats and the values shown represent mean ± SEM; ****, P < 0.0001, t-test. **(C)** Same as **(A)** but with 200 nM *MCOLN1* siRNA. ****, *P* < 0.0001, t-test with Bonferroni tests in case of samples used in multiple pairwise comparisons. **(D)** Bar graph showing fraction of T24 cells in the G_0_/G_1_ phase of the cell cycle in response to the indicated treatments for the indicated durations. Circles represent independent biological repeats and the values shown represent mean ± SEM; ***, *P* < 0.001, t-tests. **(E)** Same as **(D)** but showing BrdU-labeled cells in S phase. **, *P* < 0.01; ***, *P* < 0.001, t-tests. Drug concentrations in **(D)** and **(E)**, 200 nM *MCOLN1* siRNA and 10 μM ML-SI1. **(F)** Same as in **(A)** but in the indicated cell lines for the indicated drugs and for the indicated durations. **, *P* < 0.01; ****, *P* < 0.0001, t-tests. Concentrations, 10 μM cisplatin and 10 μM ML-SI1. Abbreviations: n.s., not significant.

Analysis of the fractions of T24 cells in various stages of cell cycle revealed that *MCOLN1* knockdown or TRPML1 inhibition led to a significant increase in the fraction of cells in the G_0_/G_1_ phase of the cell cycle (Figure 5D). Accumulation of cells in G_0_/G_1_ was evident within 2-days of treatment and persisted at the 5-day mark (Figure 5D). The larger fraction of cells in G_0_/G_1_ was accompanied with a decline in the fraction of BrdU-labeled cells that were in the S phase (S-BrdU) (Figure 5E) and G2/M phases (Figure S4B). Therefore, the decline in T24 cancer cell numbers upon the knockdown or inhibition of TRPML1 was a consequence of accumulation in the G_0_/G_1_ phase of the cell cycle with an attendant decline in the fractions in S and G_2_/M phases.

Since ML-SI1 slowed progression through the cell cycle, we did not anticipate the drug to synergize with a cytotoxic agent. Indeed, cisplatin decreased the number of T24 and RT4 cells to similar extents (Figure 5F). Treatment with a combination of ML-SI1 and cisplatin led to a further decline in T24, but not RT4 cell numbers (Figure 5F). Normalization to the effects of ML-SI1 showed that sensitivity towards cisplatin treatment was not synergistically enhanced by TRPML1 inhibition (Figure S4C). Neither did cisplatin change the expression of *MCOLN1* (Figure S4D). Thus, ML-SI1 and cisplatin exert their effects on cancer cell number via distinct pathways leading to an additive effect upon simultaneous application of both drugs.

### TRPML1-dependent cytokine production regulates cell proliferation, cell invasion, and immune microenvironment in bladder cancers

Given the induction of *MCOLN1* and inflammation in *TP53^mut^* BLCA tumors (Figure 4), we asked whether TRPML1 could be participating in cytokine gene expression. We focused our attention on genes encoding IL6 and TNFα, which are involved in various aspects of tumorigenesis (Caetano et al., 2016; Mantovani et al., 2017; Wu and Zhou, 2010). Top half of the tumors in the TCGA BLCA dataset sorted on the basis of *MCOLN1* expression also expressed significantly higher levels of *TNF* and *IL6* than did the remained of the tumors (Figure S5A). In agreement, T24 cells expressed significantly higher levels of *IL6* and *TNF* than did HT1197 and RT4 cells (Figure 6A). Knockdown of *RELA*, which encodes the p65 subunit of NF-κB (Nolan et al., 1991), partially attenuated *IL6* and *TNF* transcription (Figure 6A). Application of ML-SI1 or *MCOLN1* knockdown *completely* mitigated the expression of the two cytokines (Figures 6A). These data indicate that cytokine gene expression in p53 deficient bladder cancer cells requires functional NF-κB and TRPML1.

**FIGURE 6.**
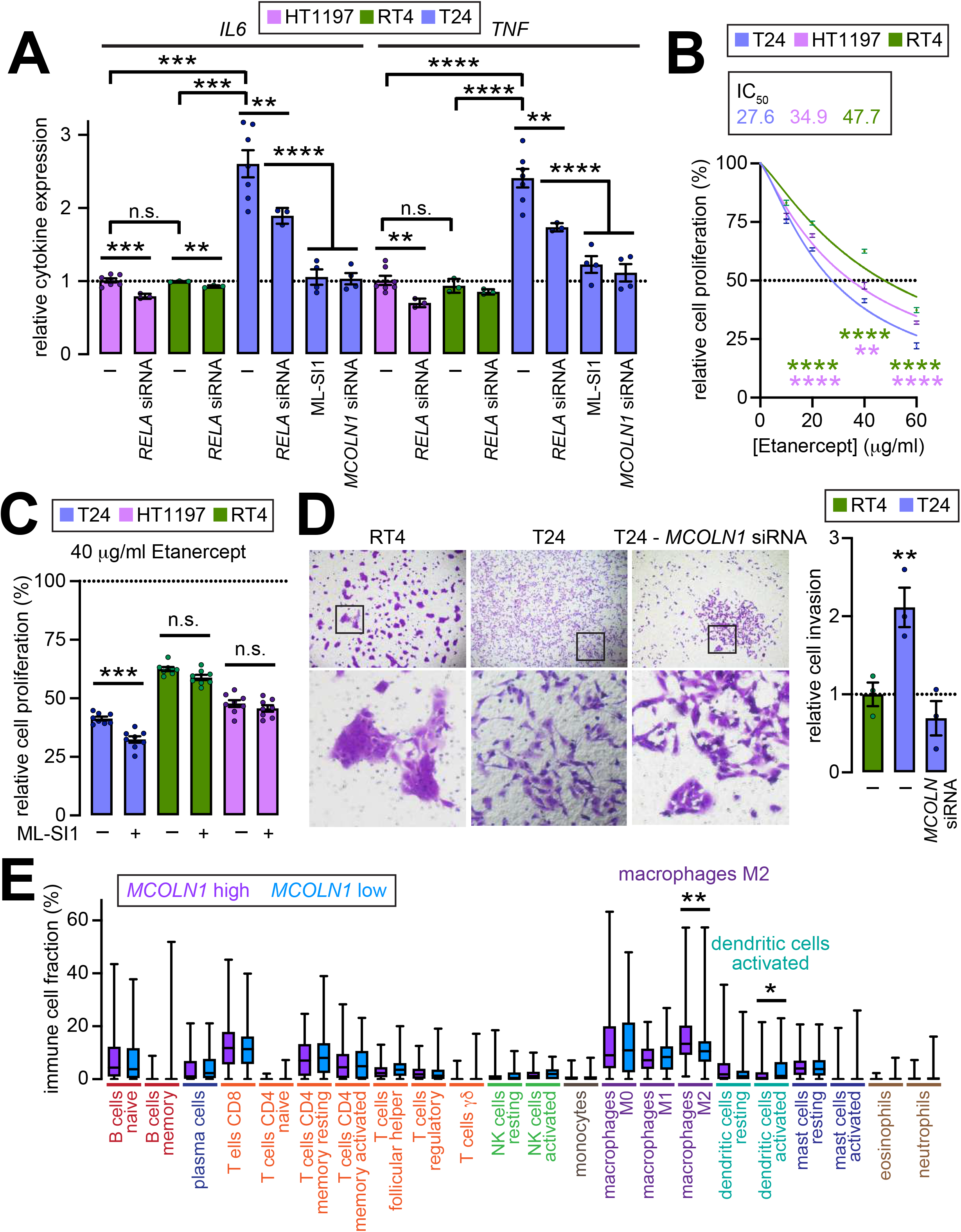
TRPML1 is required for cytokine gene expression in T24, but not HT1197, RT4, or 5637 bladder cancer cells. **(A)** Bar graph showing relative cytokine expression in the indicated cell lines treated as described. All values were normalized to the HT1197 mean. Circles represent independent biological repeats and the values shown represent mean ± SEM; **, *P* < 0.01, ***, *P* < 0.001, ****, *P* < 0.0001, t-tests with Bonferroni corrections in case of samples used in multiple pairwise comparisons. Concentrations, 50 nM *RELA* siRNA, 200 nM *MCOLN1* siRNA, and 10 μM ML-SI1. **(B)** Trace showing relative proliferation of the indicated cell lines in response to Etanercept. For each line, cell numbers were normalized to the values detected in the absence of Etanercept. Data represent mean ± SEM. Curves were obtained by fitting the data to the ‘[Inhibitor] vs. normalized response -- Variable slope’ function in Prism 8 (GraphPad), which provides the IC_50_ values shown in the box. Statistical tests involved comparison of the values in T24 cells with the values in HT1197 (magenta asterisks) and RT4 (blue asterisks) were t-tests with Bonferroni correction to account for multiple pairwise comparisons; **, *P* < 0.01, ****, *P* < 0.0001. **(C)** Bar graph showing relative cell numbers in response to 40 μg/ml Etanercept with or without ML-SI1 as indicated at the bottom. All values were normalized to the cell numbers obtained in the absence of Etanercept. Circles represent independent biological repeats and the values shown represent mean ± SEM; ***, *P* < 0.001, t-test. **(D)** *Left*, representative images of crystal violet stained cells from the indicated type in matrigel. Images in the bottom row are magnifications of the regions depicted by boxes shown in the top row. T24 cells were treated with control siRNA or 200 nM *MCOLN1* siRNA. *Right*, bar graph showing the number of invading cells relative to the RT4 mean. Circles represent independent biological repeats and the values shown represent mean ± SEM; **, *P* < 0.01, ANOVA. **(E)** Relative fractions of the indicated immune cell types in BLCA tumors sorted on the basis of *MCOLN1* expression (FPKM values). Fractions of immune cells were determined using CIBERSORT. *, FDR < 0.05, **, FDR < 0.01, Wilcoxon ranked test followed by two-stage step-up method of Benjamini, Krieger and Yakutieli. Abbreviations: n.s., not significant.

To assess the influence of TNFα on the bladder cancer cell proliferation, we applied a TNFα chelating drug, Etanercept (Suffredini et al., 1995), to T24, HT1197, and RT4 cells. Although all three lines were sensitive to Etanercept, T24 cell numbers declined by significantly greater extents than did HT1197 and RT4 (Figures 6B-6C). Simultaneous application of Etanercept and ML-SI1 induced a further decline of cell number in T24, but not RT4 or HT1197 (Figures 6B-6C). These data argue in favor of a role for *TNF*, whose expression depended on TRPML1, in promoting the proliferation of bladder cancer cells in an autocrine manner. When grown on matrigel, significantly larger number of T24 cells (TNFα high) invaded the matrix than did RT4 cells (TNFα low) (Figure 6D). Morphology of cells within the matrix also differed between the two lines. Whereas RT4 cells formed tightly packed clusters of 10-20 cells, T24 cells invaded as individuals with typical mesenchymal morphology (Figure 6D). Knockdown of *MCOLN1* in T24 cells decreased the number of invading cells, and the cells that permeated the matrix had acquired morphologies resembling those of RT4 cells i.e. elevated cell–cell association (Figure 6D). Given the established roles for TNFα in metastases (Rossi et al., 2018; Wu and Zhou, 2010; Zhu et al., 2014), our data are consistent with TRPML1 regulating the invasiveness of bladder cancer cells via the regulation of *TNF* expression.

IL6 modulates the immune microenvironment of tumors by forcing tumor associated macrophages (TAMs) to adopt the anti-inflammatory and pro-tumorigenic M2 state (Caetano et al., 2016; Chen et al., 2018; Fu et al., 2017; Mantovani et al., 2017). Using CIBERSORT (Newman et al., 2015), we found that tumors in the top half of *MCOLN1* expression harbored a significantly higher density of M2 macrophages and lower density of activated dendritic cells than did tumors with lower *MCOLN1* expression (Figures S5B and 6E). Enrichment for M2 TAMs also correlated with higher *IL6* expression (Figure S5A). Therefore, BLCA tumors with higher *MCOLN1* and *IL6* expression exhibited a pro-tumorigenic immune signature consistent with higher densities of M2 TAMs.

### TRPML1 plays a permissive, but not sufficient, role in the regulation of proliferation and inflammation in bladder cancer cells

So far, we have shown roles for TRPML1 in T24 cells in promoting proliferation, invasion, and cytokine expression. Furthermore, knockdown of wild type *TP53* in HT1197, RT4, and 5637 cells was sufficient to increase *MCOLN1* expression. These data prompted us to ask whether increased *MCOLN1* expression in *TP53* knockdown cells would be sufficient to augment proliferation and cytokine gene expression. Treatment of HT1197 and RT4 cells with *TP53* siRNA increased neither the rates of cell proliferation nor the sensitivity to ML-SI1 (Figure 7A). Similarly, *TP53* knockdown, which was sufficient to increase *MCOLN1* expression (Figure 2E), did not elevate *IL6* or *TNF* mRNA levels in HT1197 and RT4 bladder cancer cells (Figure 7B). These data demonstrate that increased *MCOLN1* expression upon loss of p53 plays a strictly permissive role in bladder cancer cell proliferation and inflammation.

**FIGURE 7.**
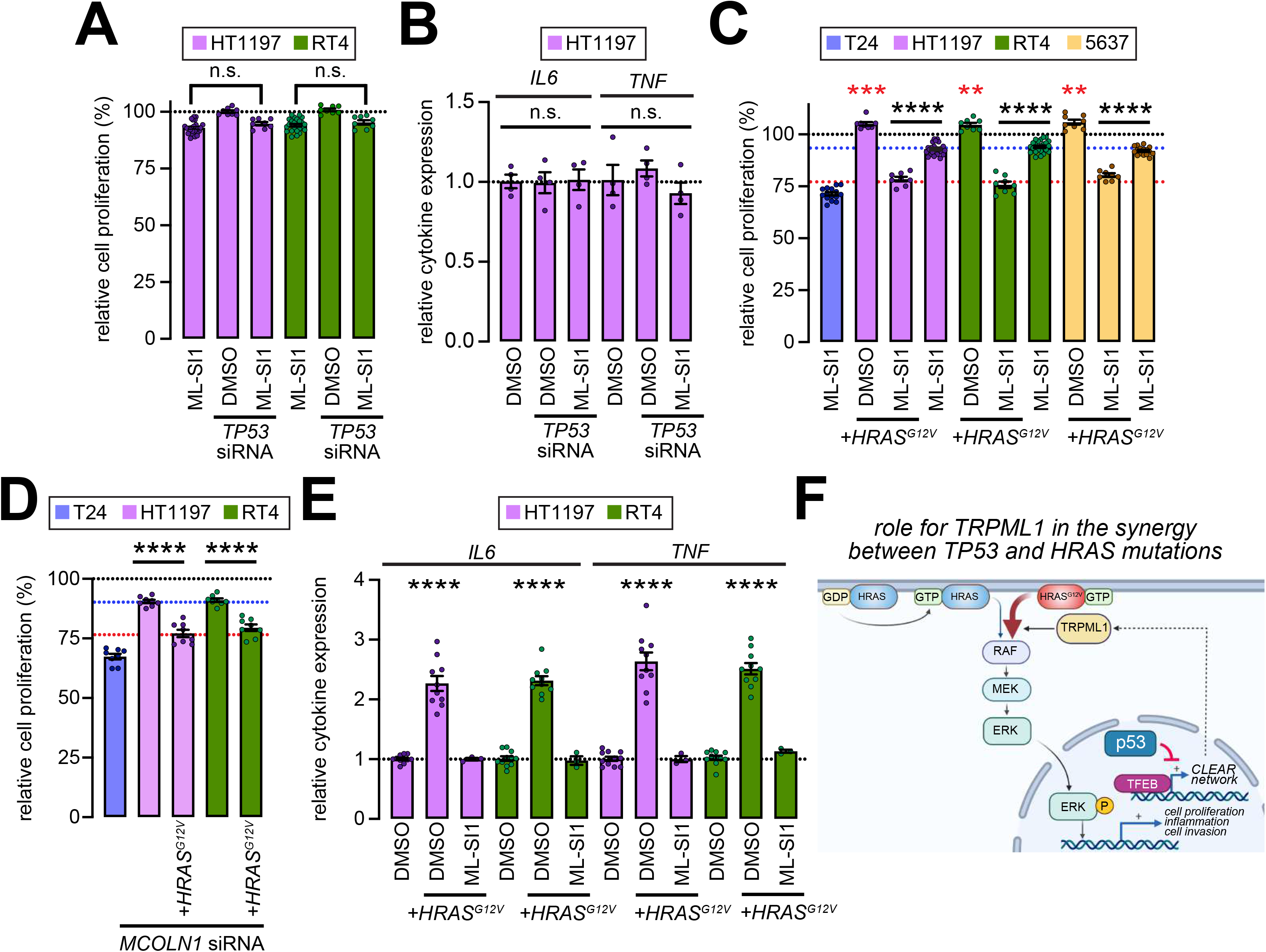
TRPML1 is necessary but not sufficient for cell proliferation and inflammation in bladder cancer cells. **(A, C, D)** Bar graphs showing relative cell numbers in the indicated lines exposed to the conditions described at the bottom. In **(A)** and **(C)**, values were normalized to HT1197 treated with DMSO alone. In **(D)**, values were normalized to HT1197 treated with control siRNA. Circles represent independent biological repeats and the values shown represent mean ± SEM; black ****, *P* < 0.0001, t-tests; red **, *P* < 0.01, ANOVA; red, ***, *P* < 0.001, ANOVA. **(B, E)** Bar graphs showing relative cytokine expression in the indicated cell lines treated as described. All values were normalized to the mean in DMSO-treated HT1197 cells. Circles represent independent biological repeats and the values shown represent mean ± SEM; ****, *P* < 0.0001, ANOVA. **(F)** Schematic showing that TRPML1 is necessary for HRAS^G12V^–MEK–ERK signaling. Therefore, increased *MCOLN1* expression in the absence of p53 permits MAPK-driven cell proliferation and inflammation. Concentrations, 75 nM *TP53* siRNA, 200 nM *MCOLN1* siRNA, and 10 μM ML-SI1. Abbreviation: n.s., not significant.

The requirement for TRPML1 in maintaining the localization and signaling-competency of HRAS^G12V^ (Jung et al., 2019) raised the possibility that bladder cancer cells lacking p53 would be highly suitable for the subsequent appearance of *HRAS* mutations. In support of this notion, lentiviral transduction of *HRAS^G12V^* into HT1197, RT4, and 5637 led to slight increases in rates of proliferation and the acquisition of *de novo* sensitivity towards TRPML1 inhibition or *MCOLN1* knockdown (Figures 7C-7D and S6A). These data indicate that oncogenic *HRAS* instilled the requirement for TRPML1 function in bladder cancer cells. Ectopic *HRAS^G12V^* also induced a 2/3-fold increase in *IL6* and *TNF* transcription (Figure 7E and S6B), which indicates that cytokine expression was driven by the MEK–ERK pathway. In agreement, inhibition of MEK1/2 using the highly-selective drug, U0126 (MEKi) (Duncia et al., 1998), attenuated expression of both *IL6* and *TNF* in T24 cells (Figure S6C). Cytokine expression driven by HRAS^G12V^ was also abolished by ML-SI1 (Figure 7E). Taken together these data indicate that increased *MCOLN1* expression following the loss of p53 has a necessary role of HRAS^G12V^-driven cell proliferation and inflammation, but is not sufficient for either in the absence of HRAS^G12V^.

## Discussion

### Correlation between *TP53* mutations and *MCOLN1* expression in bladder cancer

We demonstrate a role for p53 in regulation of *MCOLN1* expression in bladder cancer. *MCOLN1* was upregulated in primary BLCA tumors harboring *TP53* mutations predicted to ablate the transactivation function of p53, but not in tumors that were either wild type for *TP53* or harbored mutations in regions other than those that bind DNA. Given that *TP53* mutations are amongst the earliest events in bladder cancer (Gerstung et al., 2020), elevated expression of *MCOLN1* could be occurring early in the trajectory to malignancy. Conversely, increased *MCOLN1* expression in BLCA speaks to the preponderance of transactivation-deficient p53 mutations in this disease.

Ectopic knockdown of *TP53* in bladder cancer lines that do not harbor mutations in the gene was sufficient for *MCOLN1* induction. Furthermore, *MCOLN1* was constitutively overexpressed in cells that lacked p53. In only one cell line (5637 cells) was the role for p53 transactivation in *MCOLN1* repression less obvious — *MCOLN1* was not overexpressed in cells that harbored a missense mutation in the DNA binding domain (p53^R280T^). Nevertheless, ectopic knockdown of *TP53* in 5637 cells led to elevated *MCOLN1* expression. These data suggest that at least some p53 variants with mutations in the DNA binding domain retain their ability to repress *MCOLN1*, which can only be disrupted by decreasing the abundance of the mutant protein.

*MCOLN1* is under the transcriptional control of TFEB (Palmieri et al., 2011; Sardiello et al., 2009) (Figure 7F). *TFEB* knockdown mitigated *MCOLN1* upregulation in cells with constitutive absence or acute knockdown of *TP53*. We also detected an increase in *TFEB* expression upon the loss or knockdown of *TP53*. While it is possible that *TFEB* is transcriptionally repressed by p53 such that deletion or knockdown of *TP53* increases TFEB abundance, global analysis of the transcriptional targets of p53 did not point to *TFEB, TFE3* or *MITF* (Allen et al., 2014). Therefore, we anticipate hitherto unidentified p53 targets that serve as intermediaries in the repression of the TFEB–*MCOLN1* axis (Figure 7F). It is noteworthy that by phosphorylating and forcing the cytosolic retention of TFEB, TFE3, or MITF, kinases such as mTORC1, AMPK, AKT, and GSK3β regulate endolysosomal biogenesis (El-Houjeiri et al., 2019; Khaled et al., 2002; Martina and Puertollano, 2013; Martina et al., 2012, 2014; Palmieri et al., 2017; Ploper et al., 2015; Roczniak-Ferguson et al., 2012; Settembre et al., 2012). Should the activity of these kinases diminish upon the loss of p53, nuclear localization of TFEB/TFE3/MITF and *MCOLN1* expression would be increase. In agreement, loss of p53 in certain cancers leads to constitutively nuclear TFEB, and the attendant augmentation of endolysosomal biogenesis (Zhang et al., 2017). The increase in *TFEB* mRNA, then, would simply be a consequence of TFEB driving its own expression (Sardiello et al., 2009).

Our data do not necessarily argue in favor of a general role for p53 in repressing TFEB-dependent *MCOLN1* expression. Indeed, p53 activates, rather than represses, TFEB/TFE3 in normal mouse fibroblasts exposed to DNA damaging agents (Brady et al., 2018). The qualitative relationship between p53 and endolysosomal biogenesis could depend on the cell type under observation. Alternatively, inherent differences between healthy and transformed cells could determine the effects of p53 on endolysosomal biogenesis. An illuminating example of the latter is the observation that autophagosomes and endolysosomes prevent oncogenic transformation in healthy cells, but morph into tumor-promoting entities in transformed cells (Amaravadi et al., 2016; White, 2015). By extension, whether autophagy and endolysosomal biogenesis correlates with the progression or attenuation of tumorigenesis could depend on p53 status, which agrees with p53-dependent endolysosomal biogenesis opposing tumorigenesis in healthy cells (Amaravadi et al., 2016; White, 2015), whereas endolysosomal biogenesis and *MCOLN1* overexpression upon loss of p53 accelerate malignancy — the central tenet of the present study.

### TRPML1 supports oncogene-induced proliferation in p53-deficient bladder cancer cells

Increased *MCOLN1* expression in *TP53^mut^* BLCA tumors accompanied the induction of gene sets related to cell proliferation. In agreement, TRPML1 was needed for the proliferation of p53-deficient and HRAS^G12V^-positive, T24 cells. The effect TRPML1 inhibition on T24 proliferation was associated with an increased fraction of cells in the G0/G1 phase of the cell cycle, and a corresponding decrease in the fraction of cells in Sand G2/M-phases. Since G_1_ arrest is an outcome of MEK–ERK inhibition in RAS transformed cancer cells (Chambard et al., 2007; Vasjari et al., 2019; Yamamoto et al., 2006), the role for TRPML1 in HRAS^G12V^–ERK signaling that we demonstrated previously (Jung et al., 2019) explains the phase of the cell cycle that is sensitive to TRPML1 inhibition. Notably, since ML-SI1 slowed progression through the cell cycle, the drug did not synergize with the cytotoxic agent, cisplatin. Rather, simultaneous application of ML-SI1 and cisplatin led to a purely additive effect on T24 cell number.

Although *TP53* knockdown in RT4 and HT1197 cells increased *MCOLN1* expression, these alterations were not sufficient to accelerate cell proliferation. Since loss of p53, though insufficient to initiate neoplasia, predisposes cells towards accumulating oncogenic mutations (Schwitalla et al., 2013), we asked whether induction of oncogenic signaling could be more favorable in cells with higher *MCOLN1* expression. Suggesting an affirmative answer to this question, introduction of *HRAS^G12V^* heightened the sensitivity of RT4 to TRPML1 inhibition. These data suggest that *MCOLN1* induction, although not tumorigenic *per se*, sets the stage for oncogenic HRAS, which requires higher TRPML1 abundance (Figure 7F) (Jung et al., 2019). Since TRPML1 recycles and maintains plasma membrane cholesterol (Jung et al., 2019), any signaling axis that is dependent on surface cholesterol would in principle be sensitive to TRPML1 inhibition. If so, increased *MCOLN1* expression following the loss of p53 would favor proliferation driven by several oncogenic pathways.

### TRPML1 supports oncogene-induced inflammation in p53-deficient bladder cancer cells

Consistent with loss of p53 triggering inflammation (Schwitalla et al., 2013), gene sets related to inflammation were upregulated in *TP53^mut^* BLCA tumors. Likewise, *IL6* and *TNF* mRNA were significantly higher in T24 compared to RT4 or HT1197 cells. Since endolysosomal proteins such as TRPML1 are needed for NF-κB-dependent cytokine production (El-Houjeiri et al., 2019; Sun et al., 2015; Visvikis et al., 2014; Wong et al., 2017), *MCOLN1* knockdown or TRPML1 inhibition ablated the induction of cytokines. Increased *MCOLN1* expression, however, was not sufficient to augment cytokine gene expression. Rather, this requirement for TRPML1 also depended on the presence of activated HRAS, which drove *IL6* and *TNF* expression via MEK as an intermediary.

Since TNFα promotes the proliferation of malignant cells (Gakis, 2014; Ham et al., 2016; Michaud, 2007; Wang et al., 2014; Zhu et al., 2014), Etanercept diminished cancer cell numbers. T24 cells, however, exhibited greater sensitivity to Etanercept than did RT4 and HT1197 cells, which suggests a teleological explanation for TRPML1-dependent cytokine production. Indeed, Etanercept-induced decline in T24 cell number was enhanced by ML-SI1. Since TNFα also promotes cancer cell invasion (Rossi et al., 2018; Wu and Zhou, 2010; Zhu et al., 2014), T24 cells were significantly more invasive than were RT4 or HT1197 cells. *MCOLN1* knockdown mitigated the invasiveness of T24 cells, which agrees with TRPML1 being needed for *TNF* expression.

Secreted by primary cancer cells and interacting stromal entities such as fibroblasts, IL6 compels TAMs into the anti-inflammatory, M2 state (Caetano et al., 2016; Chen et al., 2018; Cho et al., 2018; Fu et al., 2017; Mantovani et al., 2017). In agreement with TRPML1 being needed for augmented *IL6* expression, BLCA tumors with higher *MCOLN1* expression exhibited significantly greater density of M2 macrophages. Given that M2 TAMs discourage the infiltration of anti-tumorigenic T lymphocytes (Mantovani et al., 2017), our data raise the possibility that higher *MCOLN1* expression is predictive of an immune-cold tumor microenvironment, and thus, poor patient prognosis (Gardner and Ruffell, 2016; Gu-Trantien et al., 2013). Future studies could evaluate whether the simultaneous application of TRPML1 inhibitor and checkpoint blockers enhance the efficacy of immunotherapy.

## Acknowledgements

We thank the Center for Advanced Microscopy, Department of Integrative Biology & Pharmacology at McGovern Medical School for the use of microscopes and cameras.

We are grateful to Drs. Guangwei Du and Dung-Fang Lee for helpful comments on the manuscript. This work was supported by the NINDS grant, R01NS081301, and NIA grant, R21AG061646 (both to K.V.).

## Author Contributions

K.V. conducted the bioinformatic analyses and J.J. performed most of the described experiments. H.L. and C.D. analyzed cell proliferation. H.L. and J.F.H. generated and provided key reagents. K.V. wrote the manuscript.

## Materials and Methods

### Bioinformatic analyses

#### Analyses of differential gene expression

Raw counts (CPM) from the RNA-seq analyses of BLCA tumors and matched controls were obtained from GDC (https://portal.gdc.cancer.gov). Differential expression was determined using EdgeR (Robinson et al., 2010) with FDR < 0.05 as the cutoff.

#### Analyses of correlation with *MCOLN1*

Spearman’s correlation coefficients of coexpression with *MCOLN1* in BLCA were obtained from cBioPortal. FDR < 0.05 was the cutoff for determining genes that were significantly correlated with *MCOLN1*.

#### GSEA

GSEA was performed using the standalone GSEA_4.0.3 application downloaded from www.gsea-msigdb.org. Pre-ranked GSEA was performed using either custom gene sets or gene sets obtained from MSigDB. For the analyses, no collapse was used, and the default minimum and maximum exclusion criteria of 15 and 500, respectively.

#### Analyses of networks

We used CytoScape to generate networks using the results of GSEA analyses. Gene sets were nodes and extent of overlap between the sets represented edge values. Node size represented number of genes, whereas node color represented normalized enrichment score (NES) from GSEA analyses.

#### Determination of Shannon information

List of somatic mutations (SNPs and small INDELs) that were determined using MuTect2 variant aggregation and masking were obtained from the UCSC Xena functional genomics browser (https://xenabrowser.net/datapages/). Using a custom R script, we filtered all synonymous mutations, and genes that did not belong to the consensus list of 722 CGCs (https://cancer.sanger.ac.uk/census; Supplementary Table 3). Tumors were ranked on the basis of *MCOLN1* FPKM values. The ranked set was then partitioned into groups comprised of tumors that belonged to either the top third or the bottom third of the ranked list. Conditional probability that mutations in a gene would occur in tumors that belonged to a group was calculated using the formula:

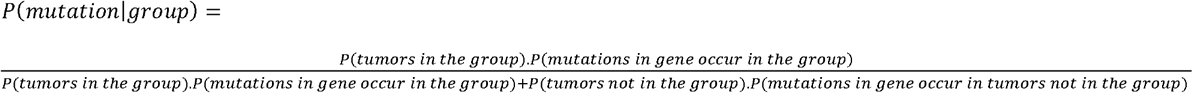

Entropy associated with gene mutations occurring in a group was calculated using the formula:

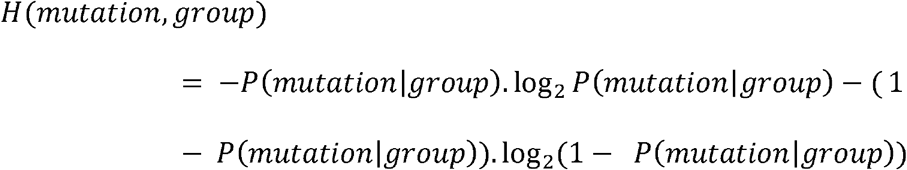

Shannon information was calculated using the formula:

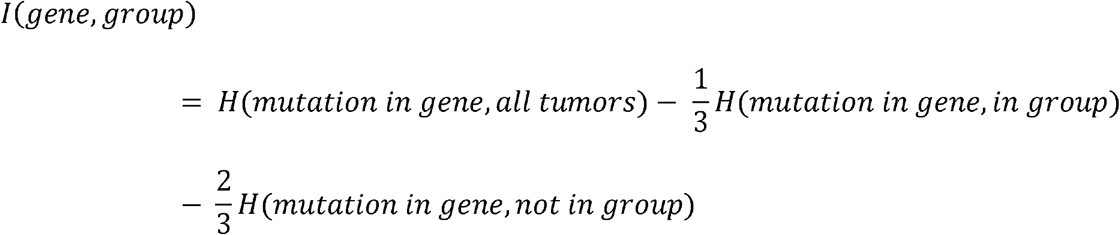

For the permutation test, we first generated a null dataset comprised of information values for each gene obtained after randomly reordering the assignment of *MCOLN1* FPKM values. Default number of permutations was 1000. Nominal *P*-values were calculated by comparing computed information for each gene with the respective values from the null dataset using a one-sample t-test. FDR *Q* values were calculated from the nominal *P*-values by applying the Benjamini and Hochberg correction. R code for running these analyses is available for download from GitHub (https://github.com/kvenkatachalam-lab/analysis-of-entropy).

#### CIBERSORT analyses

Distribution of immune cell types in the BLCA tumors was determined using the online portal (https://cibersort.stanford.edu/index.php). Fractions of each immune cell type were then compared in groups of tumors sorted on the basis of *MCOLN1* FPKM values. Pairwise comparisons were made using the Mann-Whitney test, and FDR was analyzed using the two-stage step-up method of Benjamini, Krieger and Yakutieli.

### Cell culture

T24, HT1197, RT4, 5637 cell lines (all from ATCC) were cultured in DMEM high glucose media supplemented with 10% (v/v) inactivated fetal bovine serum (FBS), penicillinstreptomycin, non-essential amino acids, pyruvate (all from Fischer Scientific), and 2mM L-glutamine (Invitrogen). All of the cell lines were maintained at 37°C and 5% CO_2_. For drug treatment, we added DMSO (vehicle) or the drugs directly to the cell media.

### Gene knockdown by RNA interference

For siRNA transfections, cells were transfected with siRNA oligonucleotides using X-treme GENE 9 DNA Transfection reagents (Roche) following manufacturer’s instructions. Subsequent analyses were performed after 2- or 5-days of siRNA treatment. The siRNA sequences were designed as described previously (Jung et al., 2019). The following sequences were used:

*MCOLN1* — 5’-CCCACATCCAGGAGTGTAA-3’ (200 nM)
*TP53* — 5’-AGACTCCAGTGGTAATCTA-3’ (75 nM)
*RELA* — 5’-GGAGTACCCTGAGGCTAT-3’ (50 nM)
*TFEB* — 5’-CCGCCTGGAGATGACCAACAA-3’ (50 nM)

Control siRNA (SIC001, Sigma-Aldrich) was used as a negative control. Concentration of control siRNA was equal to that of the specific siRNA.

### Lentiviral transduction

To ectopically express HRAS^G12V^ in bladder cancer cells, we transduced cells with a *HRAS^G12V^-GFP* lentiviral construct. Briefly, we treated cells with cell culture media containing lentivirus and 4 μg/ml of hexadimethrine bromide (Sigma) for 12 hours. After 12 hours, we removed this media and added full growth media. We incubated the cells for 48 hours before adding media with puromycin (1:5000 dilution) for selection of cells expressing *HRAS^G12V^-GFP*.

### Analyses of cell proliferation

#### WST-1 assay

Cells were seeded in 96-well plates at the density of 10^4^ cells/well and incubated for 12 hours at 37°C (CO_2_). Subsequently, fresh media containing DMSO or drugs or control or specific siRNA was applied to the cells. Cell numbers were assessed 2- or 5-days later using the WST-1 (4-(3-(4-iodophenyl)-2-(4-nitrophenyl)-2H-5-tetrazolio)-1,3-benzene disulfonate, Roche Applied Sciences) assay as per the manufacturer’s protocol. Absorbance was measured using a microplate ELISA reader (Cytation 5, BioTek) at 450 nm.

#### Cristal violet staining

Cells were seeded in 6-well plates (2×10^5^ cells/well), and exposed to 10 μM MLSI for the indicated durations. Cells were then fixed for 5 minutes with 4% PFA, and stained for 10 minutes with 0.5% crystal violet (Sigma) diluted in PBS. After a series of washes in plain water, plates were dried and images were captured. Subsequently, crystal violet signal from the plates was solubilized in 100 μl methanol, and absorbances of the solutions were read at 540 nm.

#### Gene expression analysis

When performing RT-qPCR, total RNA was extracted with RNeasy Mini Kit (Qiagen) following manufacturer’s instructions. 1μg of total RNA was reverse-transcribed with High Capacity cDNA Reverse Transcription Kit (Applied Biosystems). Real-time qPCR was performed using SYBR Green JumpStart Taq ReadyMix (Sigma) as per the manufacturer’s protocol. Primers used were as follows:

*GAPDH:*
Forward: 5’-GAAGGTGAAGGTCGGAGTC-3’
Reverse: 5’-GAAGATGGTGATGGGATTTC-3’
*IL6:*
Forward: 5’-AGACAGCCACTCACCTCTTCAG-3’
Reverse: 5’-TTCTGCCAGTGCCTCTTTGCTG-3’
*MCOLN1*:
Forward: 5’-CTGGTGGTCACGGTGCAG-3’
Reverse: 5’-CTGCTCCCGCGTGTAGG-3’
*RELA*:
Forward: 5’-TGAACCGAAACTCTGGCAGCTG-3’
Reverse: 5’-CATCAGCTTGCGAAAAGGAGCC-3’
*TFEB*:
Forward: 5’-CCAGAAGCGAGAGCTCACAGAT-3’ Reverse: 5’-TGTGATTGTCTTTCTTCTGCCG-3’ *TNFA:*
Forward: 5’-CTCTTCTGCCTGCTGCACTTTG-3’ Reverse: 5’-ATGGGCTACAGGCTTGTCACTC-3’

### Analyses of cell cycle

Number of cells in G1, S or G2/M phases of the cell cycle was determined by measuring DNA content using propidium iodide (PI) and bromodeoxyuridine (BrdU) labeling followed by flow cytometry analysis. Cells were pulsed with 10 μM BrdU for 45 minutes, harvested and fixed with 70% ice-cold ethanol overnight. Cells were washed in BrdU wash solution (0.5% Tween 20, 0.5% BSA in PBS) and resuspended in 2 N HCl for 20 minutes at room temperature. 0.1 M sodium borate was added to neutralize, and the cells were washed twice with BrdU wash solution. Cells were incubated with anti-BrdU antibody (BD 347580 mouse IgG) at room temperature for 30 minutes, washed thrice in BrdU wash solution and incubated with Alexa Fluor 488 goat anti-mouse IgG (Invitrogen) for 30 minutes at room temperature. After three washes with BrdU wash solution the cells were treated with 100 ug/ml RNase and 25 μg/ml propidium iodide for 20 minutes at 37°C. Analyses were performed using a BD LSRFortessa instrument.

### Cell invasion assay

3×10^3^ cells in 100 μL serum-free DMEM were placed into the upper chamber of a 24-well Transwell insert (8 μm pore size; Trevigen) coated with 10% of Matrigel (v/v, Corning). 500 μL complete media was added to the lower chamber. After 24 hours, cells remaining on the upper membrane were removed with cotton wool. Cells that had migrated or invaded through the membrane were stained with the crystal violet (0.5%, Sigma) after fixation with 4% PFA as described above. Stained cells were then photographed and counted using ImageJ.

---

**FIGURE S1.**
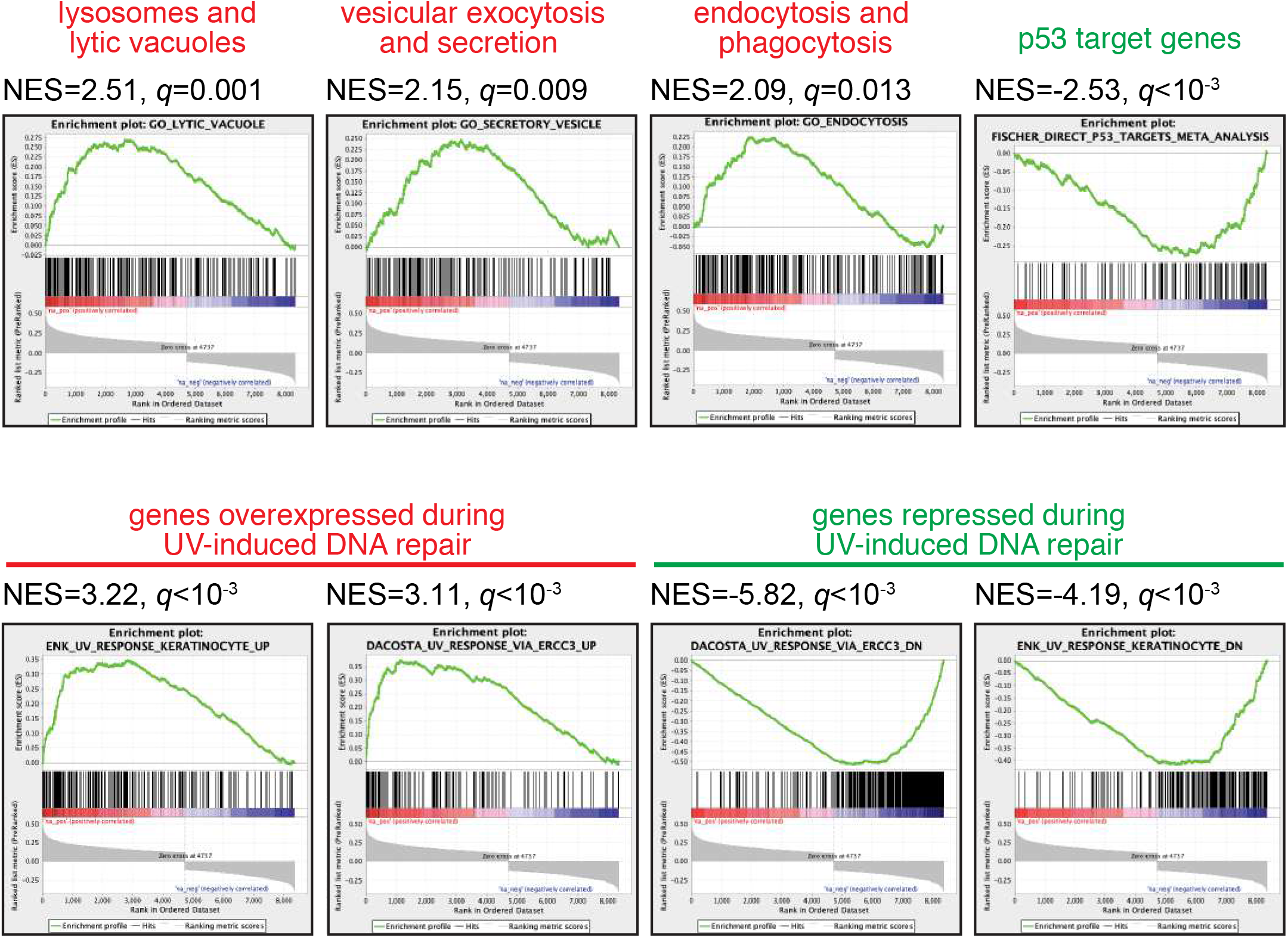
GSEA traces show enrichment for the indicated sets in genes that showed significant correlation with *MCOLN1*. Abbreviation: NES, normalized enrichment score.

**FIGURE S2.**
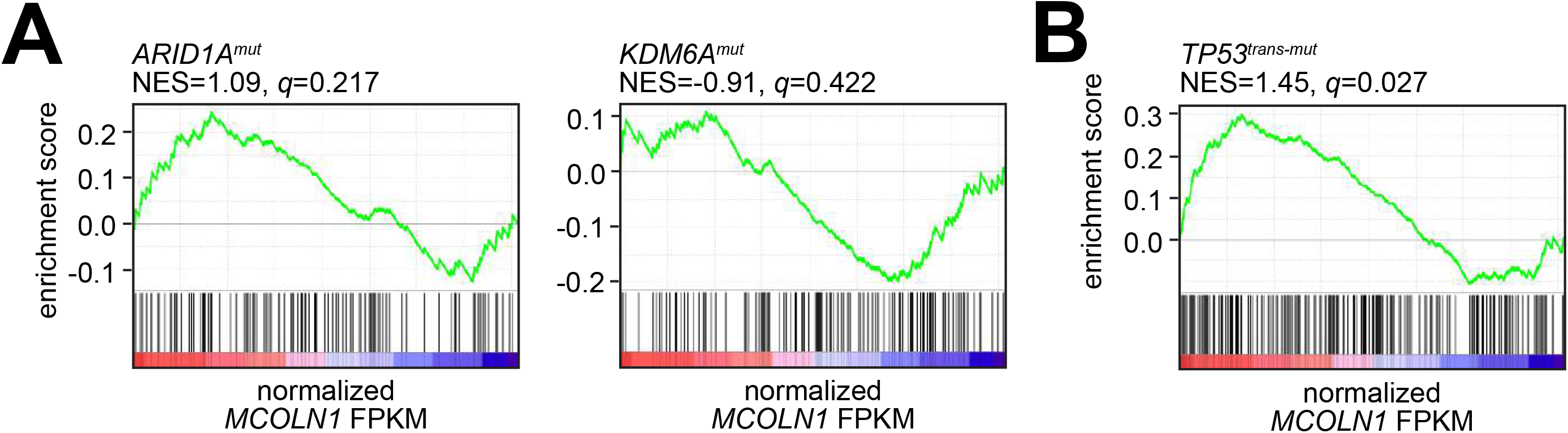
GSEA traces show enrichment for indicated mutations in tumors ranked on the basis of *MCOLN1* expression (represented as median normalized FPKM values).

**FIGURE S3.**
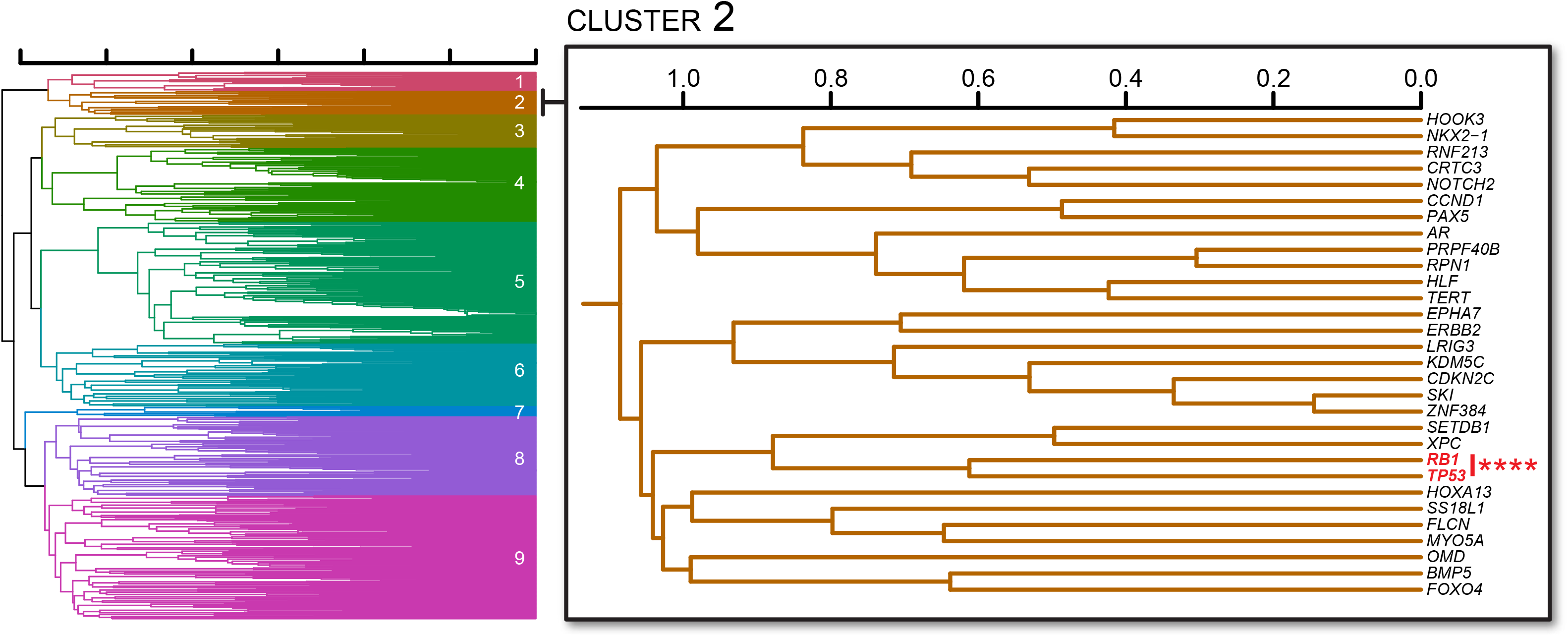
*Left*, dendrogram depicting the overlap of mutations in 723 cancer-related genes in BLCA tumors. Numbers depict 9 clusters. *Right*, magnification of cluster 2, in which *TP53* and *RB1* mutations show significant co-occurrence. ****, *P* < 0.0001, Fisher’s exact test.

**FIGURE S4.**
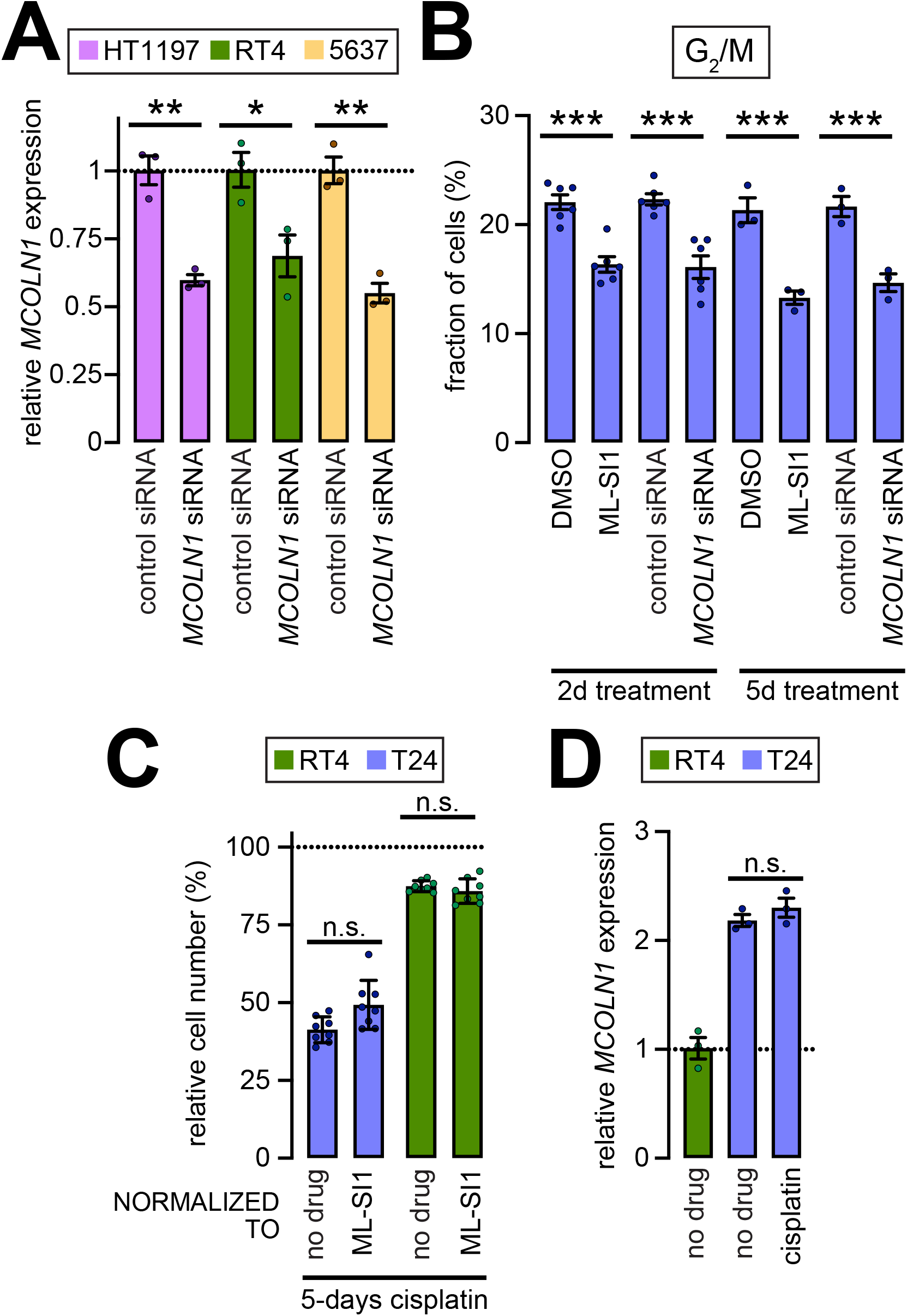
**(A)** Bar graph showing *MCOLN1* expression in the indicated cell lines treated with control or *MCOLN1* siRNA. *MCOLN1* expression was normalized to the control mean of the same cell line. Circles represent independent biological repeats and the values shown represent mean ± SEM; *, *P* < 0.05, **, *P* < 0.01, ****, *P* < 0.0001, t-tests. **(B)** Bar graph showing fraction of T24 cells in the G2/M phases of the cell cycle in response to the indicated treatments and for the indicated durations. Circles represent independent biological repeats and the values shown represent mean ± SEM; ***, *P* < 0.001, t-tests. Concentrations, 200 nM *MCOLN1* siRNA and 10 μM ML-SI1. **(C)** Relative number of cells of the indicated identities. All data were from cells treated with cisplatin for 5-days. Data used to normalize each set is mentioned at the bottom. **(D)** *MCOLN1* expression in the indicated cell lines treated as mentioned. All values were normalized to the mean in untreated RT4 cells. Circles represent independent biological repeats and the values shown represent mean ± SEM. Concentrations in **(C)** and **(D)**, 10 μM cisplatin and 10 μM ML-SI1. Abbreviation: n.s., not significant.

**FIGURE S5.**
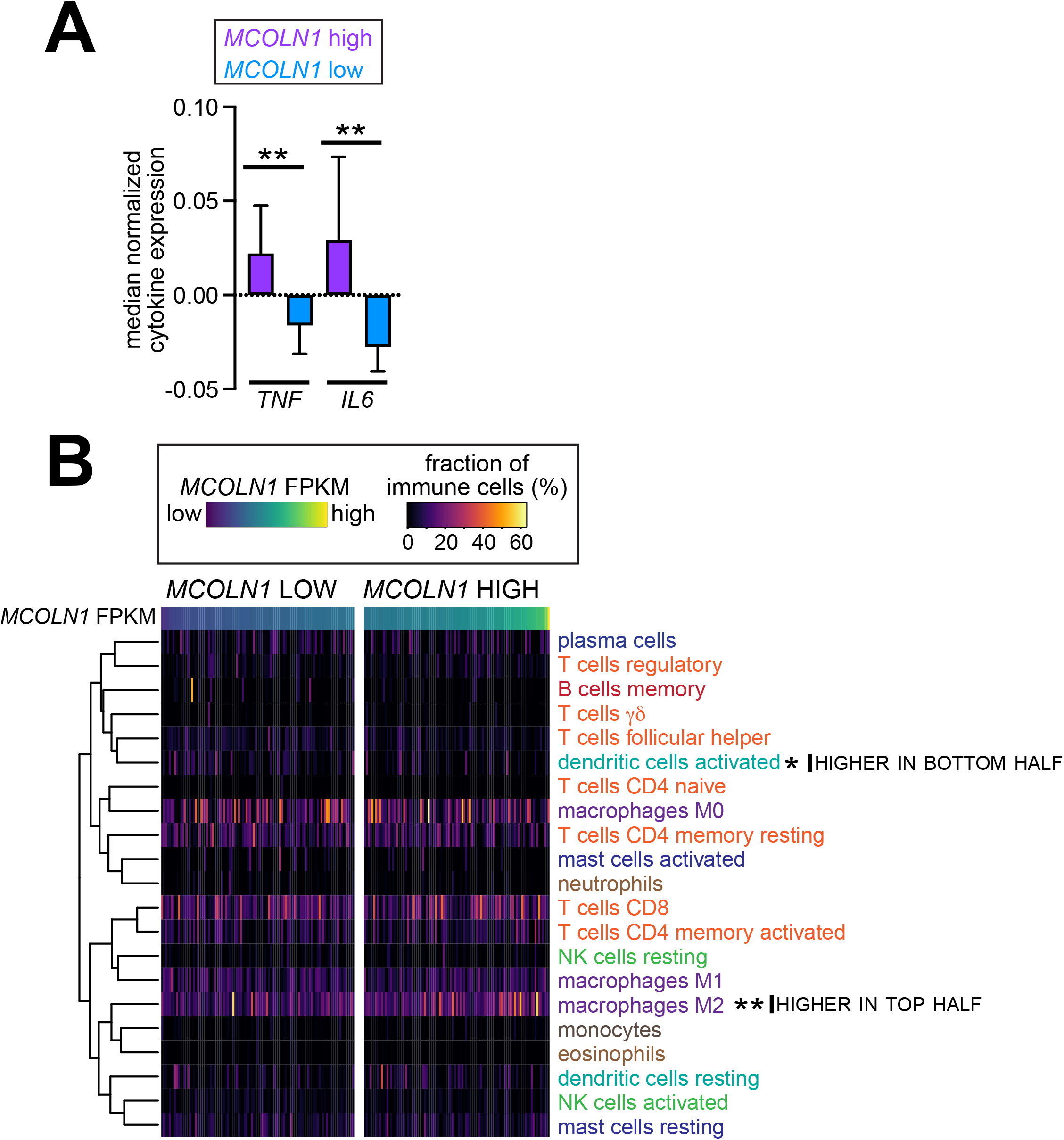
**(A)** Bar graph showing median-normalized expression of the indicated cytokines in BLCA tumors with high or low *MCOLN1* expression. Data represent mean ±95% confidence intervals. **, *P* < 0.01, Mann-Whitney test. **(B)** Heatmap depicting fractions of the indicated immune cells in BLCA tumors separated on the basis of *MCOLN1* expression. Statistical tests used are described in Figure 5E. Heatmap on the top depicts *MCOLN1* expression.

**FIGURE S6.**
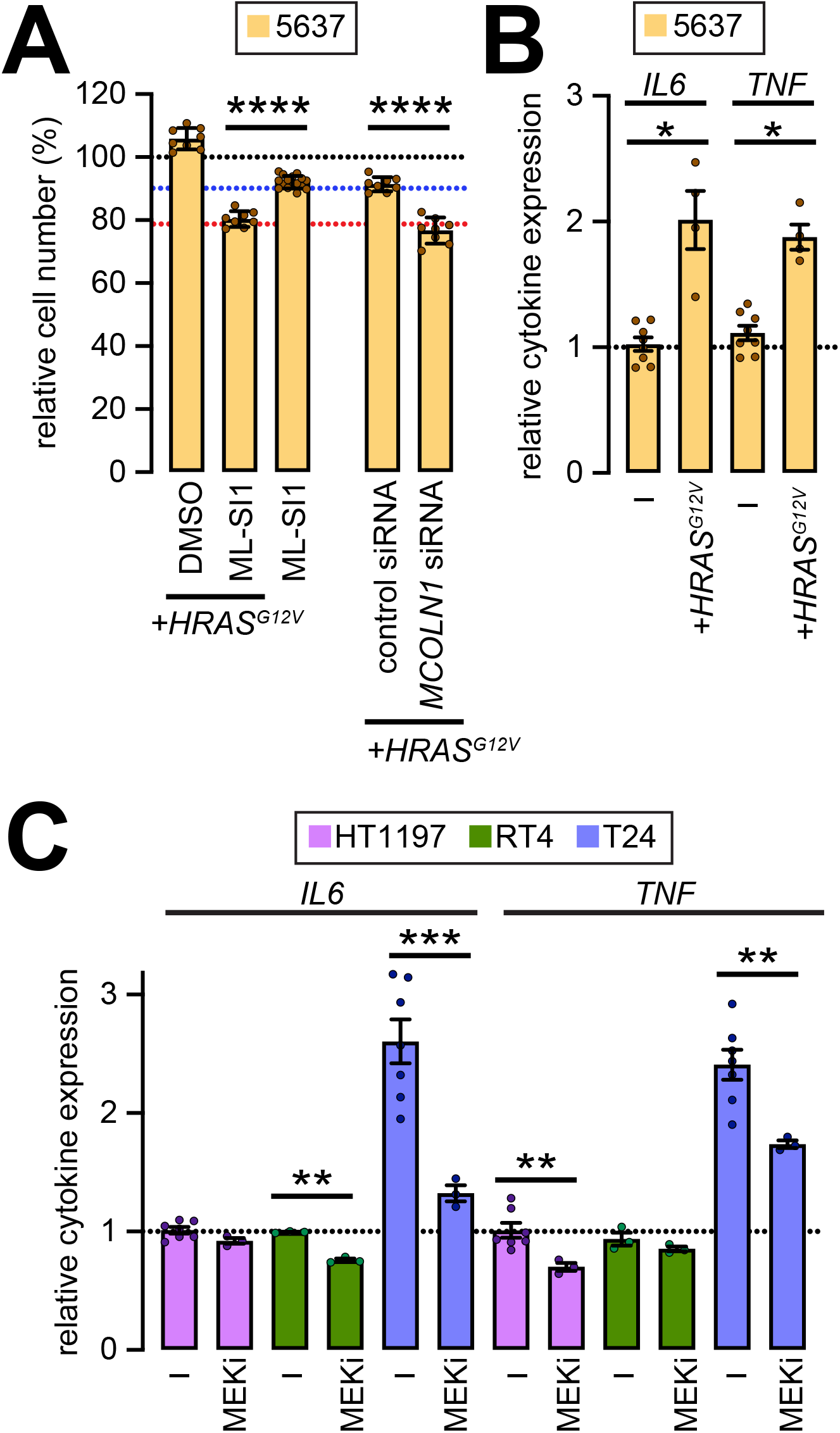
**(A)** Bar graph showing relative number of 5637 cells treated as indicated at the bottom. Values shown were normalized to untreated cells. Circles represent independent biological repeats and the values shown represent mean ± SEM; ****, P < 0.0001, t-tests. Concentrations, 200 nM *MCOLN1* siRNA and 10 μM ML-SI1. **(B)** Bar graph showing expression of the indicated cytokines in 5637 cells treated as indicated at the bottom. Values shown were normalized to untreated cells. Circles represent independent biological repeats and the values shown represent mean ± SEM; *, *P* < 0.05, t-tests. **(C)** Bar graph showing cytokine expression in the indicated cell lines treated with DMSO or 10 μM MEKi. Values shown were normalized to untreated HT1197 cells. Circles represent independent biological repeats and the values shown represent mean ± SEM; **, *P* < 0.01, ***, *P* < 0.001, t-tests with Bonferroni correction in the case of samples that were subject to multiple pairwise comparisons.

